# Botrytis virus X ORF2 suppresses RNA silencing in a trigger-restricted manner and is associated with altered Dicer-like gene expression in *Botrytis cinerea*

**DOI:** 10.64898/2026.08.20.746048

**Authors:** Fayruza Lalany, Sarah C. Drury, Mamadou L. Fall, Peter Moffett

## Abstract

RNA interference (RNAi) is a central antiviral defense mechanism in fungi, yet relatively few mycoviral suppressors of RNA silencing (VSRs) have been functionally characterized, particularly in phytopathogenic hosts. Botrytis virus X (BVX), a positive-sense RNA virus in the family *Alphaflexiviridae*, infects *Botrytis cinerea* and encodes five predicted open reading frames (ORFs), most of which have unknown functions. Here, we screened BVX ORFs 2–5 for RNA silencing suppressor activity using complementary GFP-based assays in *Nicotiana benthamiana* and examined the leading candidate in the fungal host *B. cinerea*. BVX ORF2 (X2) enhanced GFP transcript and protein accumulation in assays where silencing is triggered by sense RNA but failed to suppress silencing triggered by hairpin-derived siRNAs or miRNA-guided targeting, indicating a trigger-restricted suppressor phenotype. In *B. cinerea*, transgenic expression of X2 was associated with reduced induction of the RNAiassociated genes *BcDCL1* and *BcDCL2* compared to empty vector controls, with the strongest effect observed on *BcDCL1*. In a virus-infected fungal background, X2 expression was also associated with increased viral RNA accumulation. Together, these results identify BVX X2 as a BVX-encoded, trigger-restricted suppressor of RNA silencing and link its expression to altered RNAi-related gene induction and increased viral RNA accumulation in *B. cinerea*.

**Impact Statement:** RNA interference (RNAi) is a key antiviral defence in fungi, yet relatively few viral suppressors of RNA silencing (VSRs) have been characterized in mycoviruses, particularly in phytopathogenic hosts. This study identifies Botrytis virus X (BVX) ORF2 (X2) as a trigger-restricted suppressor that enhances GFP accumulation in plant-based assays initiated by sense RNA but does not suppress silencing initiated by defined hairpin-derived siRNA or miRNA-guided pathways. By linking this activity to altered induction of RNAi genes and increased viral RNA accumulation in *Botrytis cinerea*, this work provides one of the first experimental frameworks linking a candidate mycoviral VSR to measurable readouts of RNAi and viral RNA accumulation in a native fungal host context. These findings expand the diversity of known VSR phenotypes and establish a framework for connecting suppressor activity to RNAi-associated responses and viral outcomes in fungal systems.

## Introduction

RNA interference (RNAi) is a conserved gene regulatory mechanism that plays a central role in antiviral defense across eukaryotes (Nicolás and Garre, 2016; Ding et al., 2018; Schuster et al., 2019; Jin et al., 2021). In this pathway, Dicer-like (DCL) proteins process double-stranded RNA (dsRNA) into small interfering RNAs (siRNAs), which are incorporated into Argonaute (AGO) proteins to guide sequence-specific degradation or repression of viral transcripts (Baulcombe, 2004). In antiviral and transgene-silencing contexts, RNAi can be triggered by viral replication intermediates, highly expressed or aberrant sense transcripts that are converted into dsRNA, exogenously expressed hairpin dsRNA, or endogenous miRNA-guided targeting pathways. In fungi, RNAi-related mechanisms, including quelling, contribute to antiviral defence (Romano and Macino, 1992; Cogoni and Macino, 1999a; b; Nuss, 2011) and have been described in multiple filamentous species, including *Neurospora crassa*, *Cryphonectria parasitica*, *Fusarium graminearum*, *Magnaporthe oryzae*, *Aspergillus nidulans*, *Colletotrichum higginsianum*, *Sclerotinia sclerotiorum*, and *Botrytis cinerea* (Kadotani et al., 2004; Segers et al., 2007; Hammond et al., 2008; Campo et al., 2016; Mochama et al., 2018; Yu et al., 2020; Khalifa and Macdiarmid, 2021). Despite the central role of RNAi in fungal antiviral defence, comparatively little is known about how mycoviruses counteract RNAi pathways in their natural hosts (Segers et al., 2007; Rodriguez Coy et al., 2022).

Viral suppressors of RNA silencing (VSRs) are widely described in plant and insect viruses and act through diverse mechanisms, including sequestration of small RNAs, inhibition of Dicer activity, and interference with Argonaute function (Csorba et al., 2015). In contrast, only a limited number of mycoviral VSRs have been characterized, and reported examples suggest a relatively narrow set of phenotypes (Rodriguez Coy et al., 2022), often involving suppression of transcriptional induction of core RNAi genes such as DCL2 and AGO homologues. For example, the hypovirus protein P29 suppresses the induction of DCL2 and AGO2 in *C. parasitica* (Segers et al., 2006; Sun et al., 2009), while ORF2 of *F. graminearum* virus 1 represses DCL2 and AGO1 expression (Yu et al., 2020). More recently, Fan et al. (2024) reported a mycovirus-associated suppressor phenotype involving reduced activation of host RNA silencing genes, further supporting the view that transcriptional modulation of RNAi components is an emerging theme among characterized mycoviral VSRs. Other reported VSRs, such as VP10 of *Rosellinia necatrix* mycoreovirus 3, have been associated with altered small RNA profiles in heterologous systems, although their activity in fungal hosts remains less well defined (Yaegashi et al., 2013). Together, these studies indicate that mycoviruses can encode VSRs, but the diversity of suppressor phenotypes and their functional consequences in fungal hosts remain incompletely understood.

In fungi, RNAi pathways can exhibit functional diversification, with distinct Dicer-like proteins contributing to different RNA silencing processes, including antiviral defence, transgene silencing, and endogenous regulatory pathways. In several filamentous fungi, DCL2 has been identified as a primary mediator of antiviral RNAi (Segers et al., 2007; Neupane et al., 2019), whereas DCL1 may contribute to additional or partially overlapping functions (Mochama et al., 2018). However, this division is not universal, and in some systems, including *Botrytis cinerea*, both DCL1 and DCL2 appear to contribute to RNAi-associated responses (Wang et al., 2016; Luca et al., 2024). This raises the possibility that mycoviral suppressors may act within complex and context-dependent RNAi networks. Whether they exhibit pathway- or trigger-specific activity remains unresolved.

The phytopathogen *Botrytis cinerea* infects a wide range of crops and hosts diverse mycoviruses (Khalifa et al., 2024), yet no viral suppressors of RNA silencing have been described from viruses infecting this host. Defining how mycoviruses interact with this RNAi framework is therefore essential for understanding virus–host interactions in this pathogen. Botrytis virus X (BVX) is a positive-sense RNA virus in the family *Alphaflexiviridae* that has been reported to infect *B. cinerea*, although symptoms have not been reported in this host (Howitt et al., 2006). The BVX genome encodes a replicase, a coat protein, and three additional open reading frames (ORFs) of unknown function (Howitt et al., 2006). Although BVX shares features with plant-infecting potexviruses (Úbeda et al., 2025), it lacks the canonical triple gene block module that is typically involved in viral movement and has been linked to RNA silencing suppression in plant systems (Tilsner et al., 2012). This structural difference raises the possibility that BVX relies on distinct viral factors, making it a useful system to identify non-canonical RNA silencing suppressors potentially encoded by its additional ORFs of unknown function. Identifying whether any of these proteins function as RNA silencing suppressors is therefore critical for understanding how BVX engages with the antiviral machinery of its fungal host. Whether BVX encodes a functional suppressor of RNA silencing, and how such activity relates to RNAi-associated responses in *B. cinerea*, remain unknown.

Core RNA silencing mechanisms are conserved across eukaryotes, and heterologous GFP-based assays in *Nicotiana benthamiana* provide a practical approach to identify candidate VSRs (Lingel et al., 2005; Scholthof, 2006). We screened BVX ORFs 2–5 using complementary GFP-based assays and identified ORF2 (X2) as the primary suppressor candidate. We then examined whether X2 expression was associated with altered RNAi-related responses and viral RNA accumulation in *B. cinerea*. Together, these assays connect heterologous suppressor activity with RNAi-related and viral outcomes in the fungal host.

## Results

### BVX ORF2 suppresses RNA silencing in viral replication and transient expression assays *in planta*

Viral suppressors of RNA silencing from plant viruses are commonly identified using *Agrobacterium*-mediated transient expression assays in *Nicotiana benthamiana*, in which candidate proteins are co-expressed with GFP reporters and evaluated for their ability to inhibit RNAi and sustain GFP accumulation (Li and Wang, 2022). Under these conditions, transiently expressed GFP is normally targeted by RNAi whereas co-expression of a functional VSR sustains GFP accumulation at the transcript and protein levels.

BVX encodes a replicase (ORF1; Mtr–Hel–RdRP), a coat protein (ORF3; CP), and three additional ORFs of unknown function, including ORF2 and the 3′ overlapping ORFs ORF4 and ORF5 (Fig. 1). Because BVX lacks the canonical triple gene block found in PVX, we screened BVX ORFs 2–5 for RNA silencing suppressor activity using established GFP reporter assays *in N. benthamiana* (Figs. 2–5).

**Figure 1.**
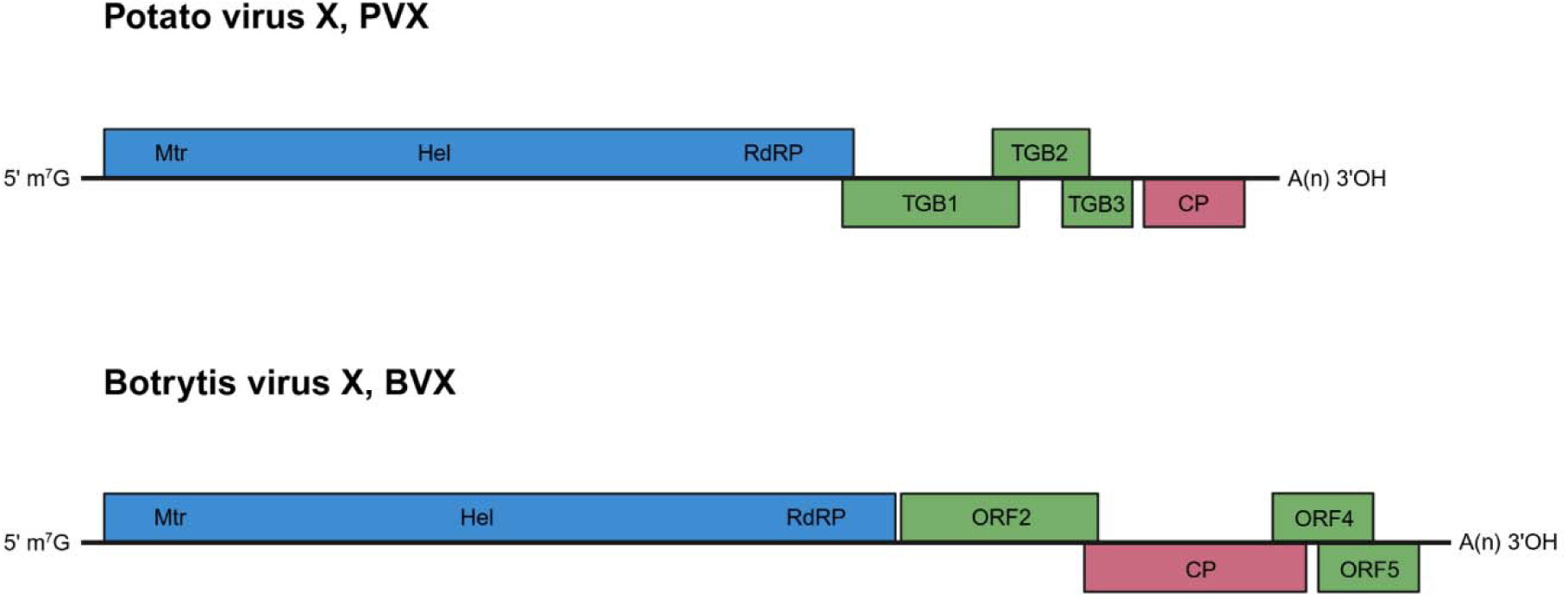
Genome organization of potato virus X and Botrytis virus X. Schematic representation of the positive-sense RNA genomes of potato virus X (PVX) (∼6.4 kb) and Botrytis virus X (BVX) (∼6.8 kb). Mtr, methyltransferase; Hel, helicase; RdRP, RNA-dependent RNA polymerase; TGB1–3, triple gene block proteins; CP, coat protein; ORF, open reading frame.

**Figure 2.**
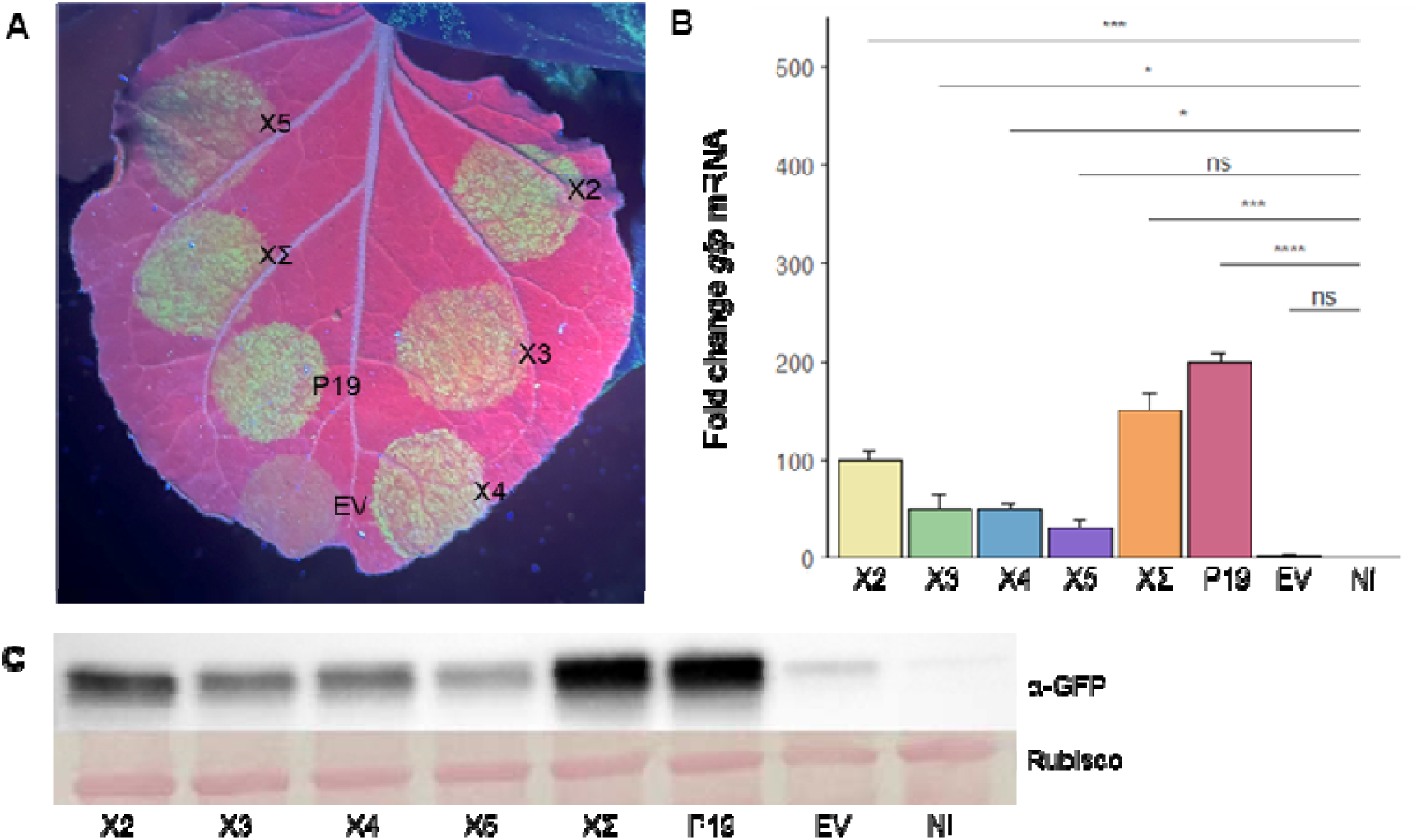
BVX ORF2 enhances GFP accumulation in the PVXΔP25–GFP replication-based silencing assay. A suppressor-deficient PVXΔP25–GFP construct was co-expressed in *Nicotiana benthamiana* leaves with BVX ORF2 (X2), ORF3 (X3), ORF4 (X4), ORF5 (X5), BVX ORFs 2–5 combined (XΣ), P19, or empty vector (EV). GFP accumulation was assessed at 5 days post-infiltration (dpi) by (A) UV illumination, (B) RT-qPCR quantification of *gfp* transcripts normalized to NbL23 and expressed relative to non-infiltrated (NI) controls, and (C) immunoblot analysis using anti-GFP antibody, with the Rubisco large subunit as a loading control. Bars represent mean ± SD (n = 9 independent plants per treatment from three independent experimental runs). Statistical analysis was performed on ΔCT values using a linear model followed by Tukey-adjusted pairwise comparisons. *P<0.05, **P<0.01, ***P<0.001, ****P<0.0001; ns, not significant.

**Figure 3.**
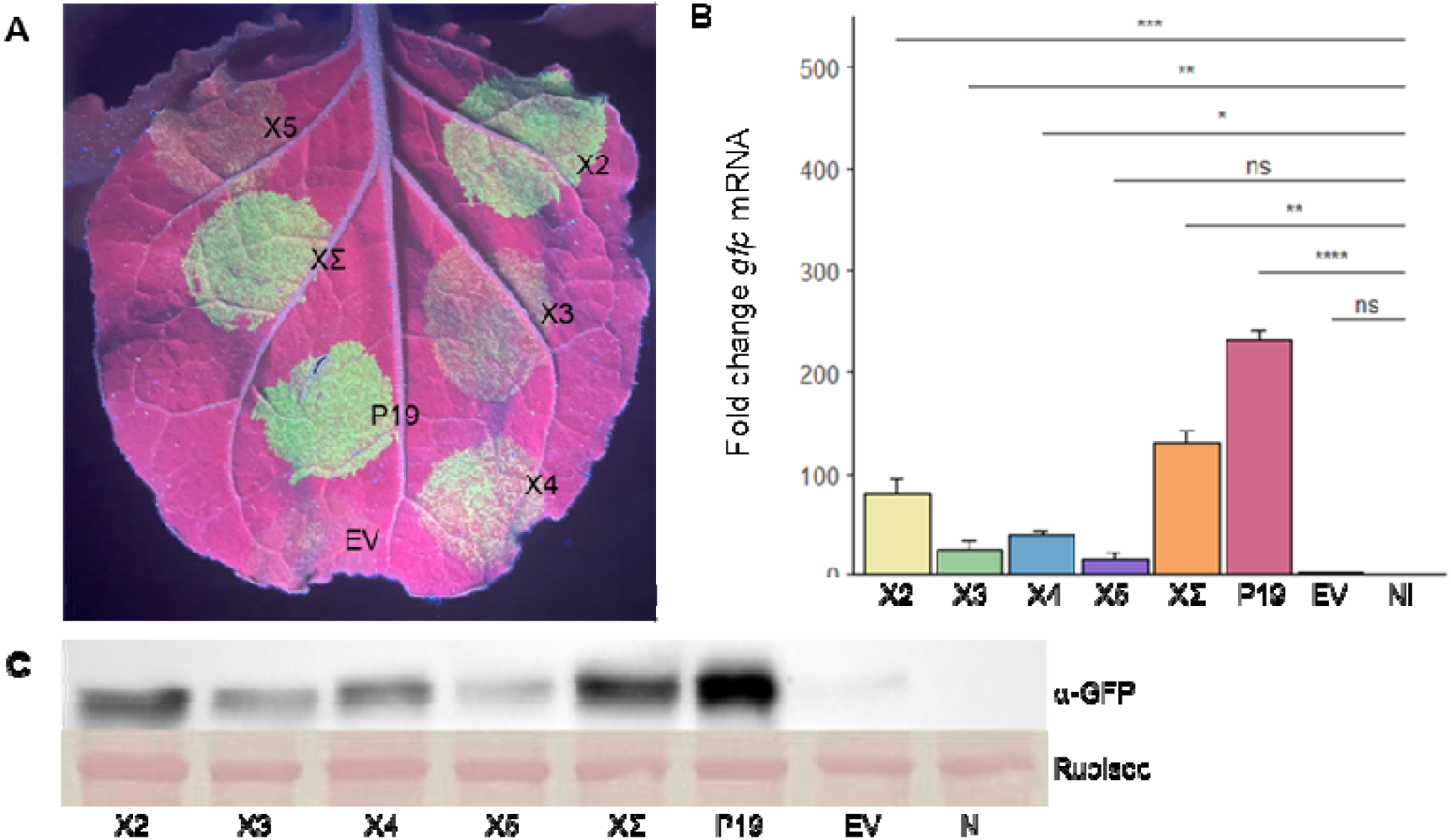
BVX ORF2 enhances GFP accumulation in a the 35S:GFP silencing assay. A 35S:GFP construct was co-expressed in *N. benthamiana* leaves with BVX ORF2 (X2), ORF3 (X3), ORF4 (X4), ORF5 (X5), BVX ORFs 2–5 combined (XΣ), P19, or empty vector (EV). GFP accumulation was assessed at 5 days post-infiltration (dpi) by (A) UV illumination, (B) RT-qPCR quantification of *gfp* transcripts normalized to NbL23 and expressed relative to non-infiltrated (NI) controls, and (C) immunoblot analysis using anti-GFP antibody, with the Rubisco large subunit as a loading control. Bars represent mean ± SD (n = 9 independent plants per treatment from three independent experimental runs). Statistical analysis was performed on ΔCT values using a linear model followed by Tukey-adjusted pairwise comparisons. *P<0.05, **P<0.01, ***P<0.001, ****P<0.0001; ns, not significant.

**Figure 4.**
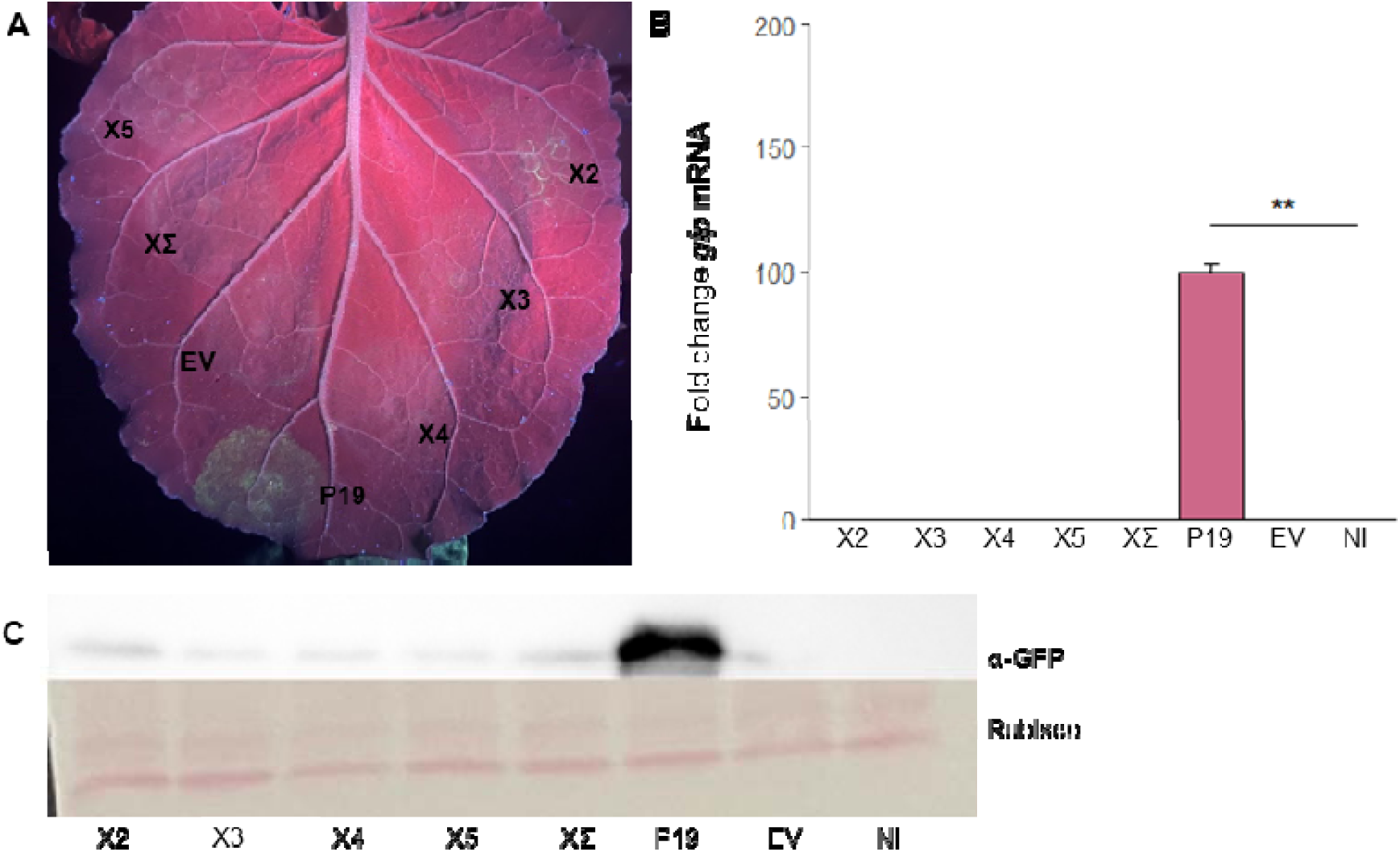
BVX ORFs do not suppress silencing triggered by hairpin-derived siRNAs. An hpGFP construct was co-expressed with 35S:GFP *in N. benthamiana* with BVX ORF2 (X2), ORF3 (X3), ORF4 (X4), ORF5 (X5), BVX ORFs 2–5 combined (XΣ), P19, or empty vector (EV). GFP accumulation was assessed at 5 days post-infiltration (dpi) by (A) UV illumination, (B) RT-qPCR quantification of *gfp* transcripts normalized to NbL23 and expressed relative to non-infiltrated (NI) controls, and (C) immunoblot analysis using anti-GFP antibody, with the Rubisco large subunit as a loading control. Bars represent mean ± SD (n = 9 independent plants per treatment from three independent experimental runs). Statistical analysis was performed on ΔCT values using a linear model followed by Tukey-adjusted pairwise comparisons. *P<0.05, **P<0.01, ***P<0.001, ****P<0.0001; ns, not significant.

**Figure 5.**
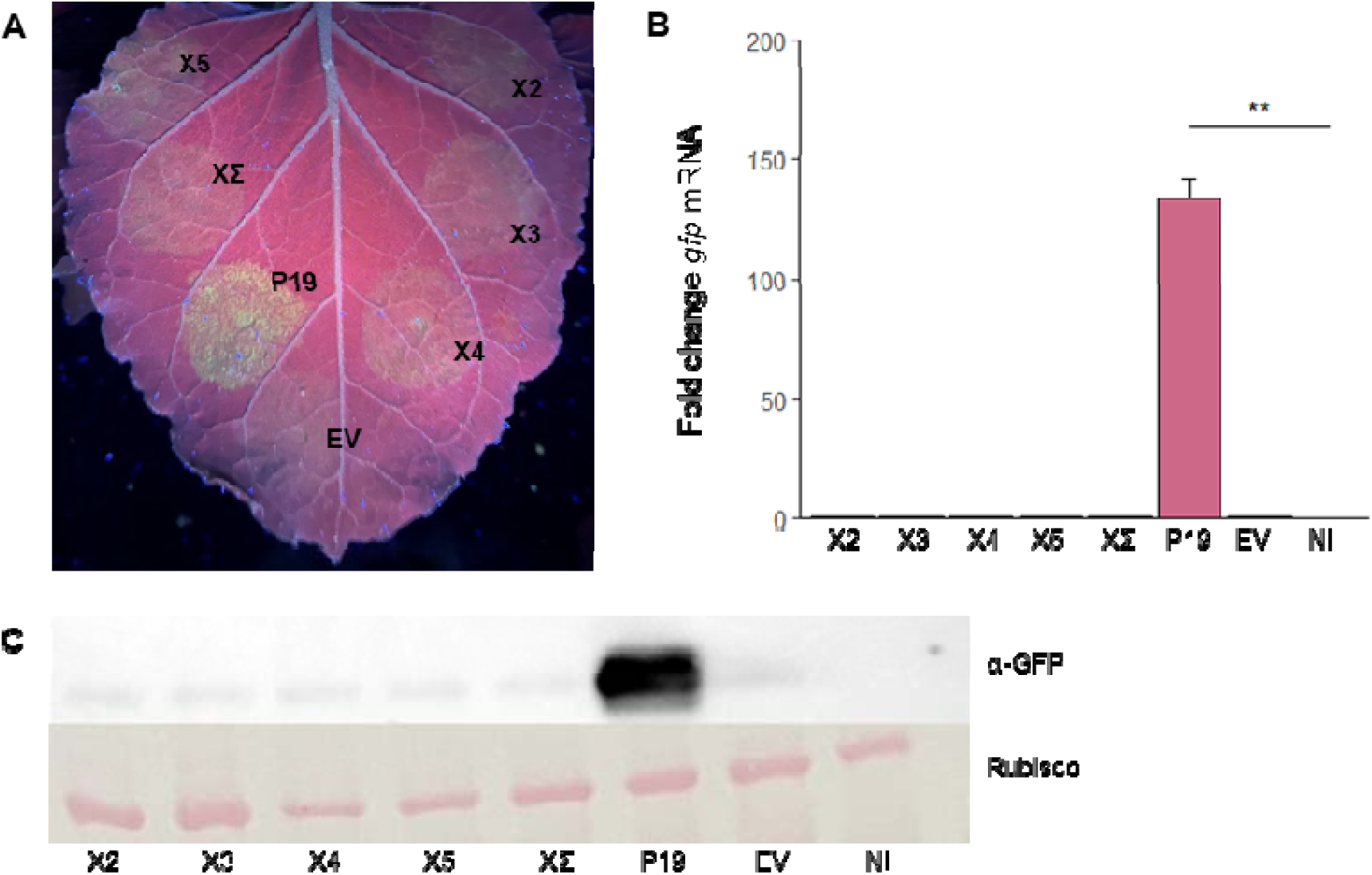
BVX ORFs do not suppress miRNA-guided silencing of GFP. A GFP171.1 miRNA reporter construct was co-expressed in *N. benthamiana*leaves with BVX ORF2 (X2), ORF3 (X3), ORF4 (X4), ORF5 (X5), BVX ORFs 2–5 combined (XΣ), P19, or empty vector (EV). GFP accumulation was assessed at 5 days post-infiltration (dpi) by (A) UV illumination, (B) RT-qPCR quantification of *gfp* transcripts normalized to NbL23 and expressed relative to non-infiltrated (NI) controls, and (C) immunoblot analysis using anti-GFP antibody, with the Rubisco large subunit as a loading control. Bars represent mean ± SD (n = 9 independent plants per treatment from three independent experimental runs). Statistical analysis was performed on ΔCT values using a linear model followed by Tukey-adjusted pairwise comparisons. *P<0.05, **P<0.01, ***P<0.001, ****P<0.0001; ns, not significant.

In the first tests, we used an assay that mimics the requirement for a VSR in viral accumulation *in planta* using a version of PVX lacking its VSR and expressing GFP (PVXΔP25–GFP), which is highly susceptible to RNA silencing. BVX ORFs (X2–5) were cloned into an expression vector for transient *Agrobacterium*-mediated expression *in planta* (materials and methods) and co-expressed individually with PVXΔP25–GFP, as well as the tombusvirus P19, a well-defined VSR positive control (Scholthof, 2006). As expected, co-expression of P19 resulted in strong GFP accumulation under UV light after five days, whereas expression with an empty vector control (EV) resulted in little visible fluorescence (Fig. 2A). Expression of X2 also resulted in strong GFP expression, as did co-expression of ORFs 2–5 (XΣ), whereas ORFs 3-5 showed an intermediate level compared to EV (Fig. 2A).

Reverse transcription quantitative PCR (RT-qPCR) confirmed a ∼200-fold increase in *gfp* mRNA in P19-expressing patches and a ∼100-fold increase in X2 patches compared to empty vector (EV) controls (Fig. 2B). ORF3 (CP; X3) and ORF4 (X4) gave intermediate effects (∼50-fold and ∼30-fold, respectively), while ORF5 (X5) showed weaker activity (∼15-fold). Co-expression of ORFs 2–5 (XΣ) yielded ∼150-fold transcript accumulation. Western blot analysis corroborated these results, with the most intense GFP bands in P19, XΣ, and X2 infiltrations, and weaker signals in X3–X5 (Fig. 2C).

Following the above results, we tested BVX proteins in a VSR assay based on transient co-expression with 35S:GFP, which measures suppression of RNAi targeting abundant transiently expressed transgene transcripts (Fig. 3). As expected, P19 enhanced *gfp* mRNA ∼230-fold and X2 ∼80-fold, compared to EV controls. X3–X5 showed only modest increases (∼25–40-fold), while XΣ produced ∼130-fold accumulation. Western blot analysis again confirmed that GFP protein levels paralleled the transcript abundance, with high expression of GFP in combination with P19, XΣ, and X2, and weaker bands in X3–X5 samples (Fig. 3C). Together, these findings identify X2 as a BVX-encoded suppressor of RNA silencing in the PVXΔP25–GFP and 35S:GFP assays.

### BVX ORFs do not suppress silencing triggered by hairpin-derived siRNAs or miRNA-guided targeting

Because X2 enhanced GFP transcript and protein accumulation in both the PVXΔP25–GFP and 35S:GFP assays, we next tested whether its activity extended to silencing initiated by defined small-RNA triggers. Specifically, we evaluated BVX ORFs in two assays that distinguish hairpin-derived siRNA silencing from endogenous miRNA-guided targeting.

In the hpGFP system, a construct expressing a hairpin of *gfp* RNA is co-infiltrated with 35S:GFP to generate abundant dsRNA, which is processed by Dicer-like enzymes into *gfp* siRNAs that efficiently and robustly silence GFP through the siRNA effector pathway. In this assay, GFP fluorescence and *gfp* mRNA were strongly reduced in empty-vector (EV) controls and robustly restored by the control VSR P19, confirming effective silencing induction and assay sensitivity (Fig. 4A–C). In contrast, none of the BVX ORFs, including X2, produced measurable rescue of GFP accumulation at either transcript or protein level (Fig. 4A–C). We next tested BVX ORFs using a GFP171.1 reporter system, which contains a GFP transgene bearing a target site for an endogenous *N. benthamiana* miRNA processed via the DCL1-dependent miRNA pathway, resulting in constitutive miRNA-guided repression of GFP (Parizotto et al., 2004). As expected, P19 increased GFP accumulation, whereas none of the BVX ORFs enhanced GFP transcript or protein levels compared to EV controls (Fig. 5A–C). RT-qPCR confirmed X2 transcript accumulation in all four assay contexts, indicating that the absence of GFP rescue in the hpGFP and GFP171.1 assays was not attributable to failure of X2 transcription (Fig. S5).

Together, these data indicate that X2 enhances GFP accumulation in the PVXΔP25–GFP and 35S:GFP assay contexts but does not measurably affect silencing initiated by hairpin-derived siRNAs or miRNA-guided targeting. This supports a trigger-restricted suppressor phenotype under the conditions tested.

### X2 expression is associated with reduced induction of RNAi core genes in *B. cinerea*

We next assessed whether X2 is associated with RNAi-related responses in its native host, *Botrytis cinerea*. A mycovirus-free isolate (BC-2016-85; see materials and methods) was transformed with constructs expressing X2, P19, or empty vector (EV), and transformants were validated by RT-qPCR and confocal microscopy based on the GFP expression cassette in pFLexpress (Figs. S1 and S2).

Transcript levels of *BcDCL1* and *BcDCL2* were quantified by RT-qPCR and normalized to an untransformed control (Fig. 6). All transformant backgrounds showed induction of both genes relative to the untransformed isolate, indicating induction of RNAi-related gene expression following transformation and transgene expression. EV transformants therefore provide a transgene-induced RNAi activation baseline. However, the magnitude of induction differed between constructs. EV and P19 strains exhibited strong upregulation, with mean increases of ∼22-fold for *BcDCL1* and ∼10.5-fold for *BcDCL2*. In contrast, X2 strains showed significantly lower induction than EV or P19, with mean increases of ∼6-fold for *BcDCL1* and ∼7-fold for *BcDCL2* (Fig. 6). These results indicate that X2 expression is associated with attenuation of RNAi gene induction relative to the transgene-induced baseline.

**Figure 6.**
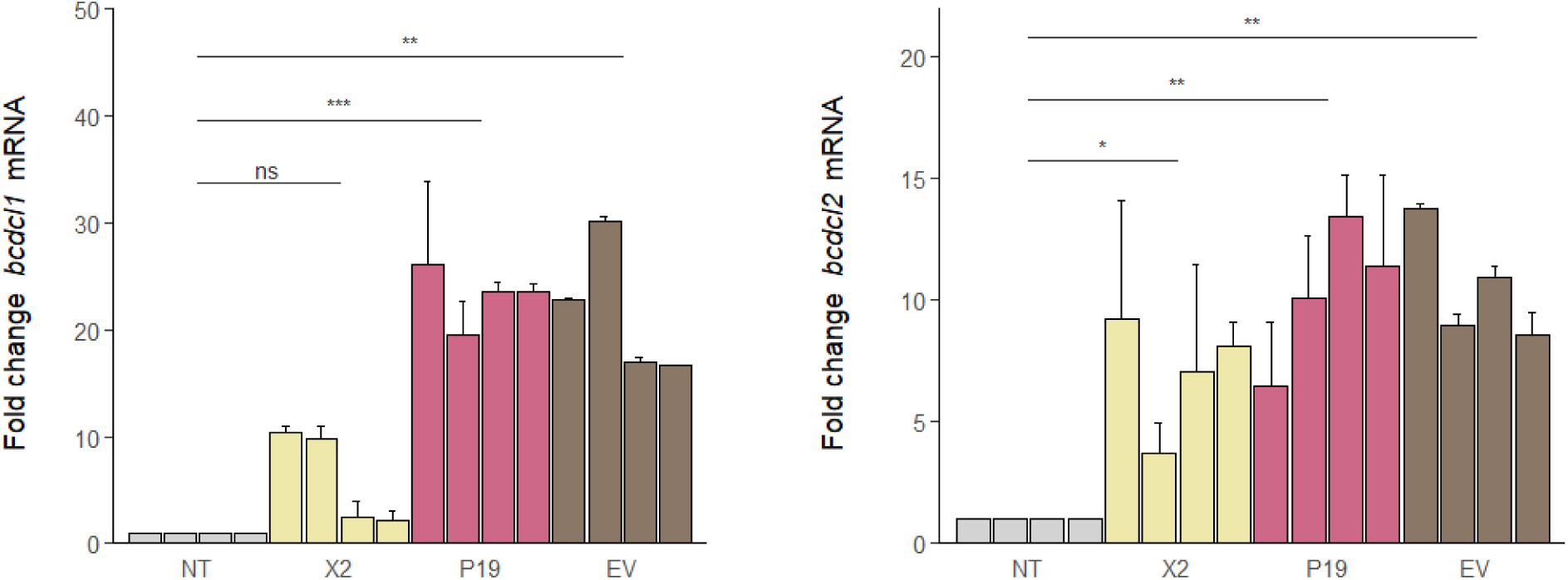
X2 expression is associated with reduced induction of RNAi-related Dicer-like genes in virus-free *Botrytis cinerea*. Relative transcript abundance of *BcDCL1* and *BcDCL2* in untransformed *B. cinerea* BC-2016-85 (NT) and transformants carrying empty vector (EV), P19, or BVX ORF2 (X2). RNA was collected at 7 days post-inoculation. Transcript abundance was normalized to 18S rRNA and expressed relative to NT; EV served as the transformed biological control. Each bar represents one independent transformant and shows the mean ± SD of three independently cultured samples. Four independent transformants were analysed per treatment ( 4). Statistical significance was assessed using a linear mixed-effects model followed by Tukey-adjusted pairwise comparisons. *P<0.05, **P<0.01, ***P<0.001, ****P<0.0001; ns, not significant.

*BcDCL1* induction was more variable among X2 transformants than *BcDCL2*, with some transformants showing modest induction and others showing near-baseline levels (Fig. 6). GFP transcript levels from the pFLexpress control cassette also varied among transformants (Fig. S2), indicating transformant-level variation in expression from the vector. In contrast, *BcDCL2* induction was more consistent across X2 transformants. Together, these data indicate that X2 expression is associated with reduced induction of *BcDCL1* and *BcDCL2* compared to the EV baseline, with the clearest reduction observed for *BcDCL1*.

### X2 expression is associated with increased viral RNA accumulation in *B. cinerea*

To test whether X2 expression was associated with viral RNA accumulation in the fungal host, we transformed a *B. cinerea* isolate (BC-2020-5) known to be infected with several viruses, including Erysiphe necator-associated fusarivirus (ENFV; see materials and methods). ENFV was selected for viral RNA quantification because it was consistently detected in BC-2020-5 and provided a robust RT-qPCR target for assessing virus accumulation.

P19- and X2-expressing transformants showed approximately 9-fold and 5-fold higher ENFV RNA levels, respectively, relative to the untransformed control, whereas EV transformants showed reduced viral RNA accumulation (Fig. 7). This pattern is consistent with increased viral RNA accumulation in suppressor-expressing backgrounds. However, the reduced ENFV levels in EV transformants indicate that transformation and/or transgene expression also influenced viral accumulation and this baseline should be considered when interpreting these data.

**Figure 7.**
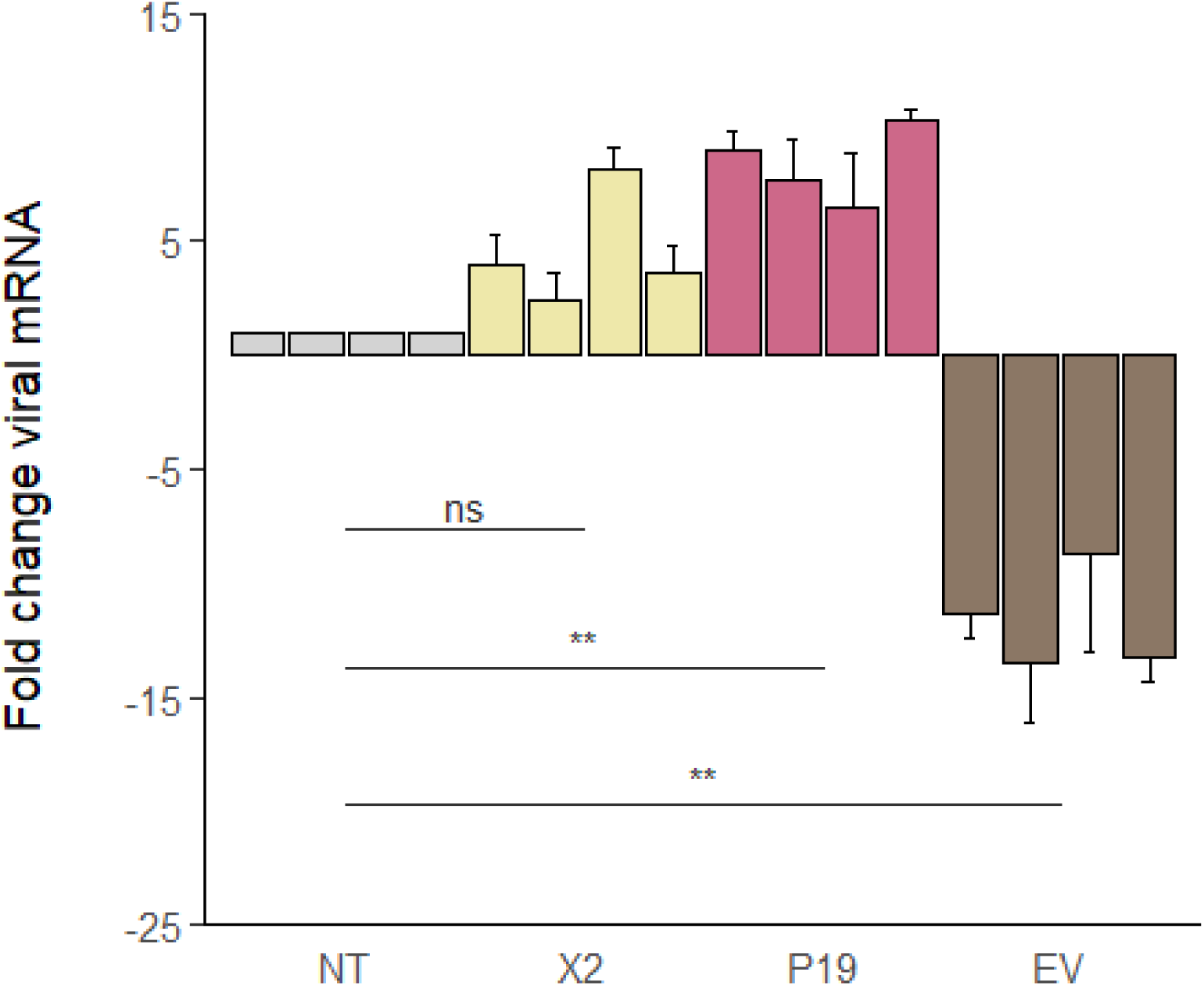
X2 expression is associated with increased ENFV RNA accumulation in ENFV-infected *Botrytis cinerea*. Relative Erysiphe necator-associated fusarivirus (ENFV) RNA abundance in untransformed *B. cinerea* BC-2020-5 (NT) and transformants carrying empty vector (EV), P19, or BVX ORF2 (X2). RNA was collected at 7 days post-inoculation. ENFV RNA abundance was normalized to 18S rRNA and expressed relative to NT; EV served as the transformed biological control. Each bar represents one independent transformant and shows the mean ± SD of three independently cultured samples. Four independent transformants were analysed per treatment ( 4). Statistical significance was assessed using a linear mixed-effects model followed by Tukey-adjusted pairwise comparisons. 0.05, 0.01, 0.001, 0.0001; ns, not significant.

## Discussion

This study identifies BVX ORF2 (X2) as a trigger-restricted suppressor of RNA silencing and links X2 expression with altered RNAi-related responses and increased viral RNA accumulation in *B. cinerea*. X2 enhanced GFP protein and transcript accumulation in the PVXΔP25–GFP and 35S:GFP silencing assays, but did not measurably suppress silencing triggered by hairpin-derived siRNAs or miRNA-guided targeting (Figs. 2–5). In *B. cinerea*, X2 expression was associated with reduced induction of *BcDCL1* and *BcDCL2* and with increased ENFV RNA accumulation in a infected-infected background (Figs. 6-7). Together, these findings identify X2 as a BVX-encoded VSR with a restricted activity profile and connect its expression to measurable RNAi-associated and viral outcomes in the fungal host.

In *N. benthamiana*, X2 reproducibly enhanced GFP accumulation in two complementary reporter systems. In the PVXΔP25–GFP assay, X2 partially restored the accumulation of a replicating viral reporter lacking its native suppressor, while in the 35S:GFP assay, X2 sustained the accumulation of an abundantly expressed transient transcript (Figs. 2–3). The concordant increases in GFP transcript and protein in both assays provide strong evidence that X2 antagonizes RNA silencing *in planta*. Its activity in the PVXΔP25–GFP system demonstrates that X2 can provide suppressor activity in a viral replication context, although this does not establish that X2 is mechanistically equivalent to PVX P25/TGB1. Because the two assays differ in reporter architecture, RNA dynamics and the processes that initiate and amplify silencing, differences in the magnitude of GFP accumulation should not be interpreted as direct differences in suppressor strength between the systems. Rather, each assay should be interpreted as a qualitative output of suppressor activity.

BVX ORFs 3–5 produced weaker effects when expressed individually, whereas pooled co-expression of ORFs 2–5 (XΣ) resulted in greater GFP accumulation than any single BVX ORF in both positive assays (Figs. 2–3). XΣ consisted of independently expressed ORFs rather than a fusion protein and did not reproduce the natural expression levels or stoichiometry of BVX proteins. The stronger XΣ phenotype therefore does not definitively establish synergy or a native BVX protein complex. It may instead reflect additive effects, altered protein dosage or combinatorial contributions from proteins that showed modest activity individually.

Nevertheless, X2 remained the strongest candidate in the single-ORF screen and was therefore selected for analysis in *B. cinerea*. The XΣ result provides evidence that other BVX proteins may contribute to suppressor-associated phenotypes, but targeted combinations and controlled expression analyses will be required to identify the contributing ORFs.

The lack of measurable X2 activity in the hpGFP and GFP171.1 assays indicates that its suppressor phenotype depends on the context in which silencing is initiated. In both assays, P19 restored GFP accumulation, demonstrating that silencing was active and that each system could detect suppressor activity (Figs. 4–5). In contrast, X2 did not increase GFP transcript or protein accumulation. X2 transcripts were detected in all four reporter contexts, reducing the likelihood that the negative results arose simply from failure of X2 transcription (Fig. S5).

The two assays in which X2 was active involve silencing initiated in the context of expressed sense RNA, whereas hpGFP supplies a defined hairpin-derived dsRNA trigger and GFP171.1 undergoes miRNA-guided targeting. This pattern supports a trigger-restricted phenotype rather than broad inhibition of downstream siRNA- or miRNA-guided silencing. One interpretation is that X2 affects a process that is particularly important during the establishment or amplification of sense-RNA-associated silencing but becomes less influential once abundant small-RNA triggers are supplied directly. Nevertheless, the experiments do not identify the molecular step targeted by X2. Small-RNA abundance, RNA-dependent RNA polymerase activity, Dicer processing and AGO-associated small RNAs were not measured. The assays therefore define the contexts in which X2 is active but do not establish whether it acts on RNAi amplification, small-RNA biogenesis, pathway regulation or another process.

The fungal experiments extend the plant-based suppressor phenotype by examining X2-associated responses in the host species of BVX. Transformation and transgene expression were associated with strong induction of both *BcDCL1* and *BcDCL2,* including in EV transformants (Fig. 6). EV therefore represents the appropriate transformed-background control for evaluating X2-associated effects. Compared to EV, X2 transformants showed reduced induction of both DCL genes, with the clearest reduction observed for *BcDCL1*. This differs from characterized mycoviral suppressors such as CHV1 P29 and FgV1 P2, which have been associated predominantly with induction of *DCL2* (Segers et al., 2006; Yu et al., 2020). The broader DCL-associated signature observed with X2 may reflect the participation of both DCL proteins in RNAi-related processes in *B. cinerea*.

The DCL results should not, however, be interpreted as evidence that X2 directly represses *BcDCL1* or *BcDCL2* transcription. Expression from the vector-borne GFP cassette varied among independently generated transformants, indicating transformant-level differences that could arise from integration site, transgene copy number, expression level or other transformation-associated effects. Differences in dsRNA load or cellular physiology may also have contributed to the DCL-expression patterns. The fungal DCL data are therefore best interpreted as an altered RNAi-associated transcriptional response linked to X2 expression rather than as proof of direct transcriptional suppression.

X2 expression was also associated with increased ENFV RNA accumulation in the infected *B. cinerea* isolate (Fig. 7). P19- and X2-expressing transformants accumulated more ENFV RNA than the untransformed isolate, whereas EV transformants showed reduced viral RNA accumulation. Because transformation itself altered ENFV abundance, EV provides the most relevant control for assessing the X2-associated phenotype. The higher ENFV levels in X2 transformants are consistent with reduced RNAi-mediated antiviral restriction, although we cannot rule out additional mechanisms. Transformation-associated changes in fungal growth, developmental state, metabolic activity or cellular fitness could also affect viral accumulation. However, the ENFV results nonetheless are consistent with an association between X2 expression and enhanced viral RNA accumulation.

The use of ENFV as a viral RNA readout was based on its consistent detection in the BC-2020-5 isolate and its performance as a robust qPCR target. Although this assay does not directly test the role of X2 during BVX infection, it provides a gain-of-function system in *B. cinerea* for determining whether X2 expression can alter the accumulation of a co-infecting virus. The ENFV result therefore complements the heterologous plant assays by linking X2 expression to a measurable viral outcome in the fungal host. Direct BVX infection or reverse-genetics systems will ultimately be required to determine the contribution of X2 to the BVX life cycle.

The potential consequences of X2 activity for fungal biology also remain unresolved. RNAi pathways in *B. cinerea* contribute to endogenous gene regulation and host-associated small-RNA processes, raising the possibility that a viral suppressor could influence fungal fitness or interactions with plants. However, the present study did not determine whether X2 expression or BVX infection alters fungal growth, morphology, pathogenicity or virulence. These phenotypes must be measured directly before conclusions can be drawn about the role of X2 during plant infection. Future studies combining BVX reverse genetics with small-RNA profiling, RNAi-deficient fungal backgrounds and infection assays will be required to connect the trigger-restricted activity of X2 to its molecular target and biological function.

In summary, BVX ORF2 exhibits reproducible RNA silencing suppressor activity in plant reporter systems involving expressed sense RNA but not in assays initiated by defined hairpin-derived siRNAs or miRNA-guided targeting. In *B. cinerea*, X2 expression is associated with reduced induction of two RNAi-related DCL genes and increased accumulation of a co-infecting viral RNA. Although the molecular target of X2 and its role during BVX infection remain unresolved, this study expands the limited range of experimentally characterized mycoviral VSR phenotypes and provides a framework for connecting heterologous suppressor activity with RNAi-associated and viral outcomes in a fungal host.

## Material and methods

### Plant material and growth conditions

*Nicotiana benthamiana* plants were grown in BM6 (Berger) substrate under a 16 h light/8 h dark photoperiod at 23 °C and ∼60% relative humidity. Light intensity was maintained at 180 µmol m ² s ¹. Plants between 4 and 5 weeks of age were used for agroinfiltration.

### Fungal isolates and culture conditions

The *B. cinerea* isolates BC-2016-85 and BC-2020-5 were obtained from the Fall laboratory (Agriculture and Agri-Food Canada, Saint-Jean-sur-Richelieu Research and Development Centre, Quebec). To verify their viral status, both isolates were analyzed by high-throughput sequencing. Isolate BC-2016-85 was screened using an updated reference database, which consistently yielded no detectable viral reads. Although we cannot categorically rule out low levels of virus, for the purpose of this study this isolate is presumed to be mycovirus-free. In contrast, BC-2020-5 was identified as co-infected with Erysiphe necator-associated fusarivirus (ENFV), *Botrytis cinerea* endornavirus 2, Botrytis porri botybirnavirus 2, Grapevine-associated botybirnavirus 1, Sclerotinia sclerotiorum dsRNA mycovirus-L, Sclerotinia sclerotiorum dsRNA mycovirus-L-WX1, and Sclerotinia sclerotiorum dsRNA mycovirus-L-WX2 (Drury et al., 2026). Isolates and transformants were cultured on potato dextrose agar (PDA) at room temperature and subcultured as required. Antibiotics were added as appropriate for selection and maintenance of transformants.

### Plasmid construction

The coding sequence of Botrytis virus X (BVX; GenBank accession NC_005132.1) was synthesized (Integrated DNA Technologies, Coralville, IA, USA). Each predicted ORF was amplified using gene-specific primers listed in Table S1.

For transient expression in *N. benthamiana*, ORFs were cloned into the binary vector pBIN61 (Bendahmane et al., 2000), which drives expression from the cauliflower mosaic virus 35S promoter, yielding constructs pBIN61-X2, pBIN61-X3, pBIN61-X4, and pBIN61-X5. Previously described constructs included pBIN61-P19, PVXΔP25–GFP, 35S:GFP, and GFP171.1 (Dalmay et al., 2000; Parizotto et al., 2004; Schwab et al., 2006).

For fungal transformation, X2 and P19 sequences were inserted into the pFLexpress fungal expression vector (Lalany and Moffett, 2025) between the *PoliC* and *Ttub* promoters, generating pFLexpress-X2 and pFLexpress-P19.

### Transient expression assays

*Agrobacterium tumefaciens*-mediated transient expression (AMT) assays in *N. benthamiana* were performed as previously described (Moffett, 2011). Briefly, binary expression constructs were introduced into *A. tumefaciens* strain C58C1 carrying the pCH32 helper plasmid. Cultures were grown to exponential phase in Luria-Bertani (LB) medium with antibiotics at 28 °C, then resuspended in 10 mM MgCl to an optical density at 600 nm (OD_600_) of 0.2.

For each assay, suspensions of a BVX ORF construct (or control construct) and the corresponding GFP reporter construct, each adjusted to OD_600_=0.2, were mixed at a 1:1 volume ratio immediately prior to infiltration. Infiltration was performed on the abaxial surface of fully expanded leaves using a needleless syringe. For co-expression of multiple BVX ORFs (XΣ), separate *Agrobacterium* cultures carrying pBIN61-X2, pBIN61-X3, pBIN61-X4, and pBIN61-X5 were pooled such that their cumulative OD600 was 0.2 prior to mixing 1:1 with the GFP reporter culture. Thus, XΣ represents pooled co-expression of independently expressed BVX ORFs rather than a fused multi-ORF product or a reconstruction of native BVX expression stoichiometry.

Four GFP-based silencing systems were used to assess RNA silencing suppression: (i) PVXΔP25–GFP, which evaluates suppression in a viral replication context; (ii) 35S:GFP, which evaluates suppression targeting abundant transiently expressed GFP transcripts; (iii) hpGFP, which induces silencing through hairpin-derived dsRNA; and (iv) GFP171.1, which reports miRNA-guided targeting of GFP. Empty pBIN61 vector (EV) served as the negative control and pBIN61-P19 served as the positive control.

GFP fluorescence was visualized under UV illumination at 3–5 days post-infiltration (dpi). For molecular analyses, 1-cm leaf discs were collected at the indicated time point, flash-frozen in liquid nitrogen, and stored at −80 °C until use. Each assay was performed in three independent experimental runs conducted at different times, using independent plants. For each treatment, three infiltrated leaves were sampled per experimental run, yielding n = 9 infiltrated leaf samples per treatment.

### Transformation of *Botrytis cinerea*

*Agrobacterium*-mediated transformation of *B. cinerea* was performed as described previously (Lalany and Moffett, 2025). *A. tumefaciens* C58C1 carrying pFLexpress constructs and pCH32 was co-cultivated with *B. cinerea* mycelia under the conditions described in Lalany and Moffett (2025). Transformants were selected on PDA supplemented with hygromycin (250 µg/mL). Resistant colonies were subcultured for three successive generations under selection.

Transformation and transgene expression were confirmed by confocal microscopy, based on GFP fluorescence from the pFLexpress cassette, and by RT-qPCR of *gfp*. Four independent transformants were generated for each construct (EV, P19, and X2). For downstream analyses, three biological replicate samples were collected from each transformant (n = 12 samples per treatment). For statistical analysis, the four independent transformants per treatment were treated as the primary experimental units, with biological replicates nested within transformants.

### Confocal microscopy

Mycelial cultures were harvested during exponential growth and mounted in phosphate-buffered saline (PBS). GFP fluorescence was imaged with an Olympus FV3000 laser scanning confocal microscope using a 40× oil objective and 488 nm excitation. Brightfield and GFP channels were acquired sequentially to prevent bleed-through. Acquisition settings were kept constant across samples within each experiment, and image processing was performed with the Olympus FV3000 native software.

### RNA extraction and cDNA synthesis

For fungal qPCR assays, mycelial tissue was harvested seven days after transfer of actively growing mycelial inoculum to fresh PDA; this time point is referred to as 7 days post-inoculation (dpi). Approximately 100 mg of infiltrated leaf tissue or fungal mycelium was flash-frozen in liquid nitrogen and ground to a fine powder. Equivalent amounts of fungal tissue were processed for each sample, and equal amounts of total RNA were used for cDNA synthesis. Total RNA was extracted using TRIzol reagent (Invitrogen, Thermo Fisher Scientific) and purified with the RNeasy Mini Kit (Qiagen, Hilden, Germany), including on-column DNase I digestion (Qiagen). RNA integrity was evaluated by agarose gel electrophoresis and purity was assessed spectrophotometrically. Only samples with A /A ratios of 1.8–2.0 and A /A ratios of 2.0–2.2 were used for downstream analyses.

First-strand cDNA was synthesized from 1 µg total RNA using 5× All-In-One RT Master Mix (Applied Biological Materials, Richmond, BC, Canada). Resulting cDNA was diluted 1:5 prior to qPCR.

### Quantitative PCR (qPCR)

qPCR was performed using a CFX96 Real-Time PCR Detection System (Bio-Rad) and SYBR Green Master Mix (Université de Sherbrooke Protein Purification Service). Each 10 µL reaction contained 4 µL diluted cDNA, 0.5 µM each forward and reverse primer, 2× SYBR Green Master Mix, and nuclease-free water. Primer sequences are provided in Table S1. Cycling conditions were 98 °C for 2 min, followed by 40 cycles of 98 °C for 2 s and 60 °C for 5 s, with a final melt curve from 65 °C to 95 °C in 0.5 °C increments.

Gene expression was normalized to *B. cinerea* 18S rRNA for fungal samples or *N. benthamiana* L23 for plant samples using the 2 ΔΔCT method (Livak and Schmittgen, 2001). Non-transformed fungal samples and non-infiltrated plant samples served as the respective calibrators. Non-transformed fungal samples and non-infiltrated plant samples served as the respective calibrators. All reactions were run in technical triplicate and averaged prior to downstream analysis.

### Protein extraction

Frozen leaf discs (1 cm diameter) were ground to a fine powder in liquid nitrogen using a pre-cooled mortar and pestle. The powdered tissue was resuspended in ice-cold extraction buffer composed of 20 mM Tris-HCl (pH 7.4), 150 mM NaCl, 1% NP-40 (Cat. No. 74385; Sigma-Aldrich, St. Louis, MO, USA), and 1 mM EDTA (Cat. No. AM9260G; Thermo Fisher Scientific, Waltham, MA, USA), supplemented with a protease inhibitor cocktail (Cat. No. 11836170001; Roche Diagnostics, Mannheim, Germany). Homogenates were incubated on ice for 30 min with occasional mixing, then centrifuged at 16,000 × g for 10 min at 4 °C to remove cellular debris. Supernatants were collected and used immediately or stored at −80 °C.

### SDS-PAGE and immunoblot analysis

Protein extracts were mixed 1:1 with 2× Laemmli sample buffer (Cat. No. 1610737; Bio-Rad, Hercules, CA, USA) and denatured at 95 °C for 5 min. Samples were briefly centrifuged and resolved on 10.5% (w/v) SDS–polyacrylamide gels. Proteins were transferred to polyvinylidene fluoride (PVDF) membranes (Cat. No. IPVH00010; MilliporeSigma, Burlington, MA, USA) using wet transfer at 100 V for 1 h at 4 °C. Membranes were blocked with 5% (w/v) non-fat dry milk in TBST (Tris-buffered saline containing 0.1% Tween-20; Cat. No. P9416; Sigma-Aldrich, St. Louis, MO, USA) for 1 h at room temperature. Membranes were then incubated overnight at 4 °C with an HRP-conjugated anti-GFP monoclonal antibody (Cat. No. sc-9996 HRP; Santa Cruz Biotechnology, Dallas, TX, USA) diluted 1:3000 in 5% (w/v) non-fat dry milk in TBST. Following incubation, membranes were washed three times for 5 min each in TBST at room temperature. Signal was detected using Clarity™ Western ECL substrate (Cat. No. 1705061; Bio-Rad, Hercules, CA, USA) and imaged using a ChemiDoc™ imaging system (Cat. No. 17001402; Bio-Rad, Hercules, CA, USA).

### Statistical analysis and data visualization

All analyses were performed in R v4.3.0 (R Core Team, Vienna, Austria). For fungal transformation assays, treatment effects were assessed using linear mixed-effects models fitted to ΔCT values, implemented in lme4 (Bates et al., 2015), with treatment specified as a fixed effect and transformant identity included as random intercept. Subcultures were treated as repeated observations nested within transformants. The four independently generated transformants per treatment were treated as the independent experimental units (n = 4 biological replicates per treatment). Post hoc pairwise contrasts were performed with Tukey adjustment using emmeans (Searle et al., 2025).

Plant qPCR data were analyzed separately for each reporter assay using linear models fitted to ΔCT values, with treatment and experimental run included as fixed effects. Treatment means adjusted for experimental run were compared using Tukey-adjusted pairwise contrasts. Each assay comprised three independent experimental runs, with three independent plants per treatment per run (n = 9 biological replicates per treatment). Relative transcript abundance was calculated using the 2 ΔΔCT method for graphical presentation.

## Author Contributions

Fayruza Lalany: Conceptualization, Data curation, Formal analysis, Investigation, Methodology, Project administration, Supervision, Validation, Visualization, Writing – original draft, Writing – review & editing.

Peter Moffett: Conceptualization, Funding acquisition, Methodology, Project administration, Resources, Supervision, Validation, Writing – review & editing.

Sarah C. Drury: Resources, Writing – review & editing.

Mamadou L. Fall: Resources, Writing – review & editing.

