## Supplementary material for "Botrytis virus X ORF2 suppresses RNA silencing in a trigger-restricted manner and is associated with altered Dicer-like gene expression in *Botrytis cinerea*": Fig. S5

### Supplementary figures

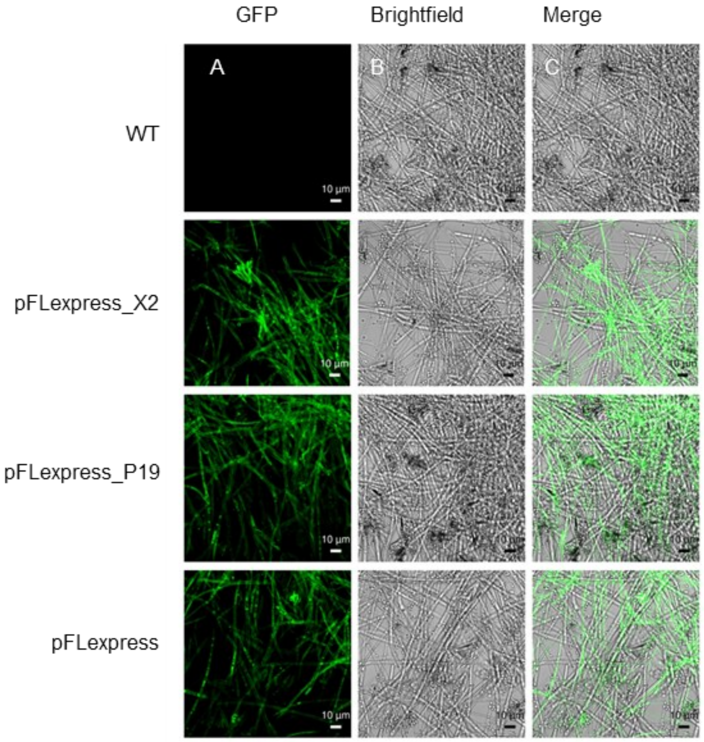


**Figure S1. GFP fluorescence in virus-free *Botrytis cinerea* transformants.**

Confocal images of hyphae from the untransformed *B. cinerea* BC-2016-85 (NT) and transformants carrying empty vector (EV), P19, or BVX ORF2 (X2). Images show (A) GFP fluorescence, (B) bright-field, and (C) merged channels. Mycelia were imaged 7 days post-inoculation. GFP was excited at 488 nm, and images were acquired using identical settings across samples. Images are representative of four independent transformants per construct and corresponding non-transformed controls. Scale bars = 10 μm.

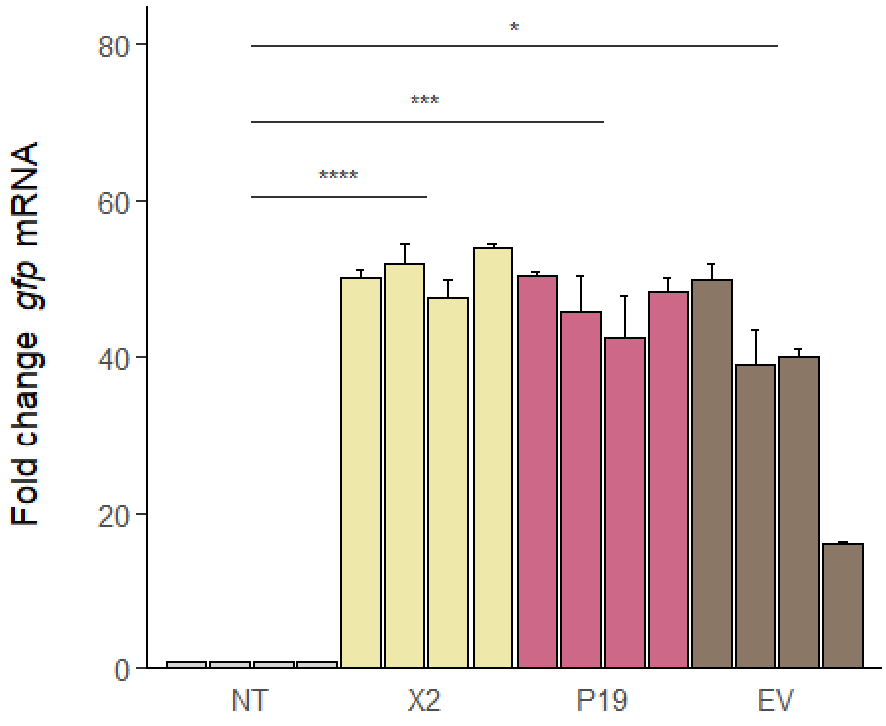


**Figure S2.** ***gfp*** **expression in virus-free transgenic** ***Botrytis cinerea*****.**

Relative *gfp* transcript levels in untransformed *B. cinerea* BC-2016-85 (NT) and transformants carrying empty vector (EV), P19, or BVX ORF2 (X2). RNA was collected at 7 days post-inoculation. Transcript abundance was normalized to 18S rRNA and expressed relative to NT; EV served as the transformed biological control. Bars show mean ± SD from four independent transformants per treatment (n=4); three independently cultured samples were analysed per transformant. Statistical significance was assessed using a linear mixed-effects model followed by Tukey-adjusted pairwise comparisons. *P<0.05, **P<0.01, ***P<0.001, ****P<0.0001; ns, not significant.

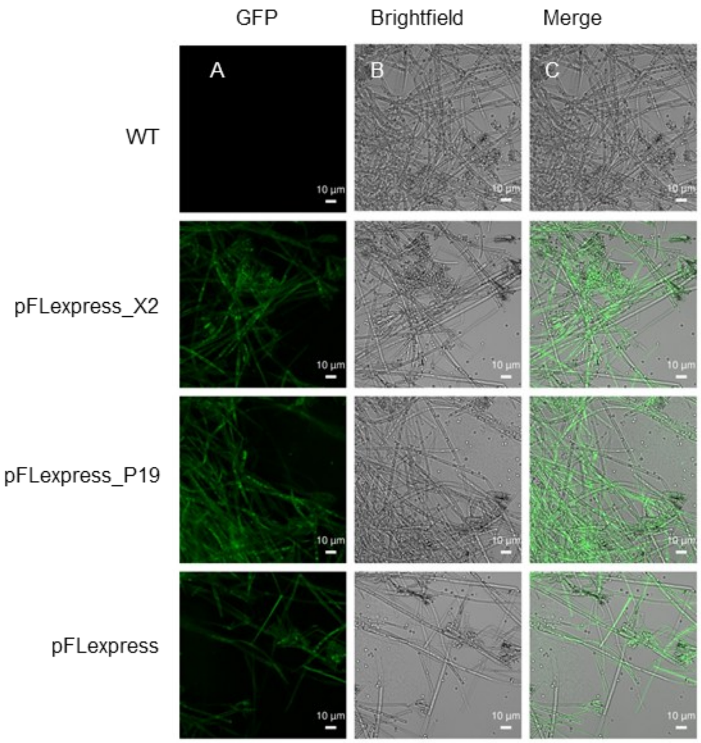

**Figure S3. GFP fluorescence in ENFV-infected *Botrytis cinerea* transformants.**

Confocal images of hyphae from the untransformed *B. cinerea* BC-2020-5 (NT) and transformants carrying empty vector (EV), P19, or BVX ORF2 (X2). Images show (A) GFP fluorescence, (B) bright-field, and (C) merged channels. Mycelia were imaged 7 days post-inoculation. GFP was excited at 488 nm, and images were acquired using identical settings across samples. Images are representative of four independent transformants per construct and corresponding non-transformed controls. Scale bars = 10 μm.

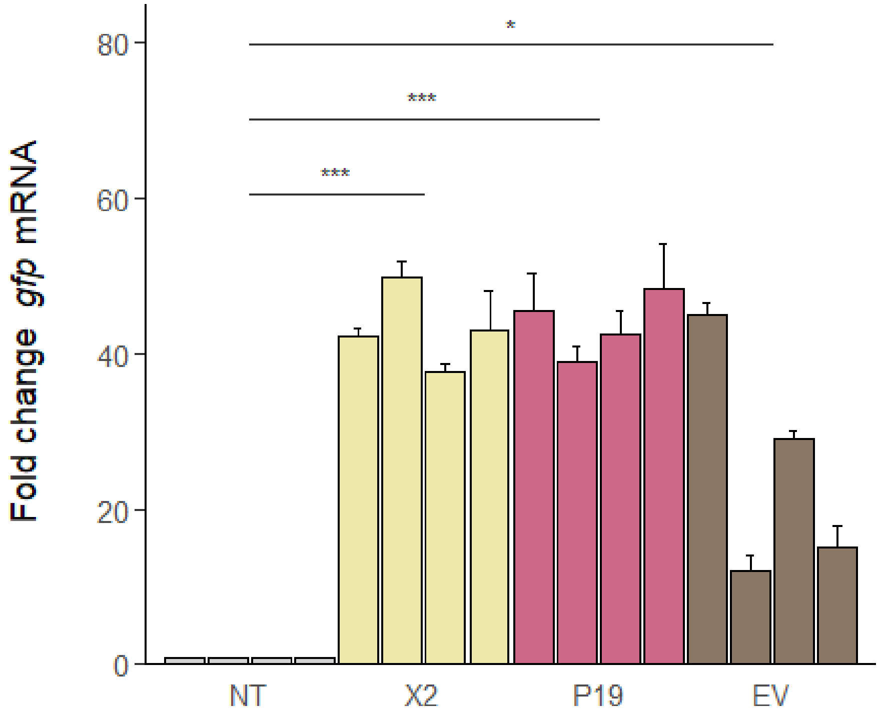


**Figure S4. *gfp* expression in ENFV-infected transgenic *Botrytis cinerea*.**

Relative *gfp* transcript levels in untransformed *B. cinerea* BC-2020-5 (NT) and transformants carrying empty vector (EV), P19, or BVX ORF2 (X2). RNA was collected at 7 days post-inoculation. ENFV RNA abundance was normalized to 18S rRNA and expressed relative to NT; EV served as the transformed biological control. Bars show mean ± SD from four independent transformants per treatment ($n = 4$); three independently cultured samples were analysed per transformant.

Statistical significance was assessed using a linear mixed-effects model followed by Tukey-adjusted pairwise comparisons. $*\, P < 0.05$, $**\, P < 0.01$, $***\, P < 0.001$, $****\, P < 0.0001$; ns, not significant.

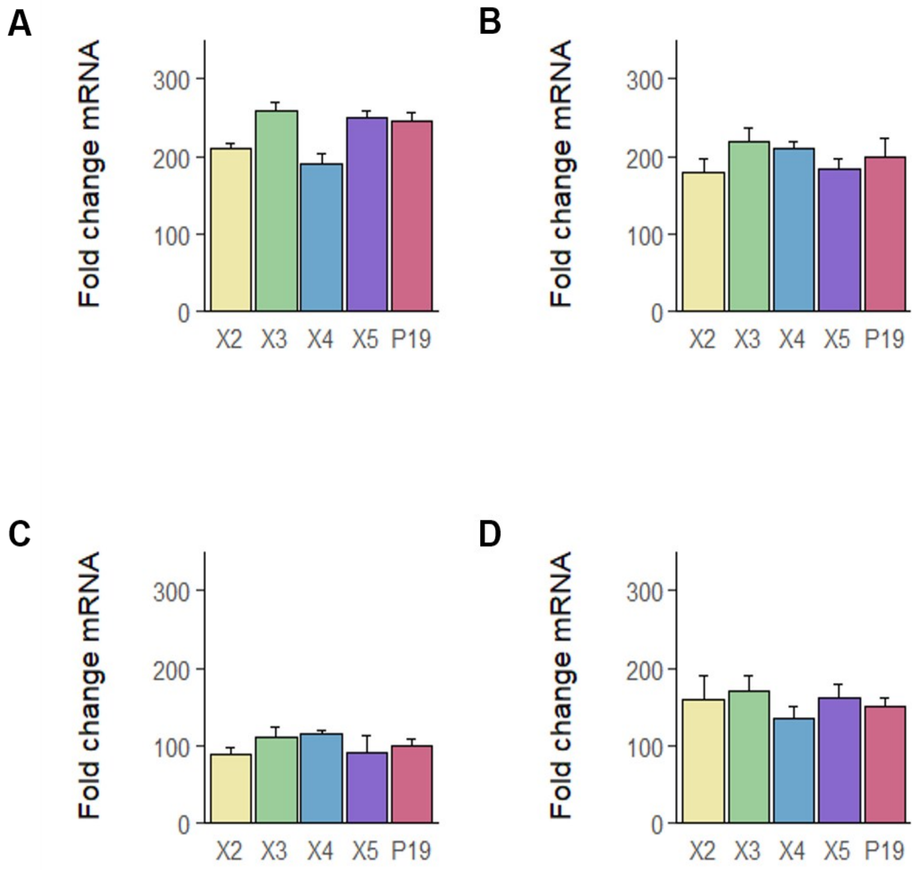


**Figure S5. Transcript accumulation of BVX ORFs and P19 across four GFP-based RNA silencing assays.**

*Nicotiana benthamiana* leaves were co-infiltrated with constructs expressing BVX ORF2 (X2), ORF3 (X3), ORF4 (X4), ORF5 (X5), or P19 together with the different reporters: (A) PVXΔP25–GFP, (B) 35S:GFP, (C) hpGFP, and (D) GFP171.1. RNA was collected 3 days post-infiltration (dpi) and transcripts were measured by RT-qPCR, normalized to *NbL23*, and expressed relative to non-infiltrated (NI) controls. Bars represent mean ± SD (n = 9 independent plants per treatment from three independent experimental runs).

### Supplementary table

**Table S1. Primers used in this study.**

Gene-specific primers used for amplification and quantitative PCR analysis of *Botrytis virus X* ORFs, GFP reporter constructs, and *Botrytis cinerea* RNAi-related genes are listed. Primer sequences are shown in the 5′–3′ orientation. Forward and reverse primers are indicated by F and R, respectively.

| Target | Primer name | Sequence (5′–3′) | Application |
|---|---|---|---|
| P19 | pFLexpress_P19.F | CGGCTCCACAGCTGCAAATGGAACGAGCTATACAAGGAAAC | Cloning into pFLexpress |
| | pFLexpress_P19.R | CTGTCTGGATCCGGTACCAAGCGATCTCTATAGCCCCCC | |

| Target | Primer name | Sequence (5′–3′) | Application |
|---|---|---|---|
| BVX ORF2 | pFLexpress_X2.F | CGGCTCCACAGCTGCAAATGTCCGTCACGCCTGAT | Cloning into pFLexpress |
| | pFLexpress_X2.R | CTGTCTGGATCCGGTACCATTAACTAATCTCAGTTAGTTGAAG | |
| NbL23 | qL23_F | AAAGCTGATCCGTCCAAAAA | RT-qPCR reference gene |
| | qL23_R | GACAGCCTTGGCAACCTTAG | |
| GFP | qGFP_F | TCCATGCCATGTGTAATCCC | RT–qPCR target |
| | qGFP_R | CCATTACCTGTCCACACAATCT | |
| 18S rRNA | q18S_F | CACCAGGTCCAGACACAATAAG | RT-qPCR reference gene |
| | q18S_R | CACTCCACCAACTAAGAACGG | |
| BcDCL1 | qBcDCL1_F | TCCATCTCCGTCTTCTTTCTCG | RT–qPCR target |
| | qBcDCL1_R | AGGGAGTGCATGAGCTACAA | |
| BcDCL2 | qBcDCL2_F | TATGGTTTCTTGCGCCTACC | RT–qPCR target |
| | qBcDCL2_R | CCACACCATCAGCACCAATA | |
| ENFV | qENFV_F | CGCGAAACCCATCAGTCTATTTCG | RT–qPCR target |
| | qENFV_R | GGCAATGAAAAGACCTTTAGAGCGG | |

| Target | Primer name | Sequence (5′–3′) | Application |
|---|---|---|---|
| BVX ORF2 | pBIN61_X2.F | TATATTCTAGAGCCACCATGTCCGTCACGCCTGAT | Cloning into pBIN61 |
| | pBIN61_X2.R | TATATGGATCCGCCACCTTAACTAATCTCAGTTAGTTGAAGTG | |
| BVX ORF3 | pBIN61_X3.F | TATATGGATCCGCCACCATGGATCCGAATTTAGATCAGGACAC | Cloning into pBIN61 |
| | pBIN61_X3.R | TATATCCCGGGGCCACCCTACTCCCCCCATCCAATCTGTG | |
| BVX ORF4 | pBIN61_X4.F | TATATTCTAGAGCCACCATGCCCACTTACTCATCAATGC | Cloning into pBIN61 |
| | pBIN61_X4.R | TATATGGATCCGCCACCCTACAGCGAGTACATCAGTGC | |
| BVX ORF5 | pBIN61_X5.F | TATATTCTAGAGCCACCATGTCCGACTTCAACGAC | Cloning into pBIN61 |
| | pBIN61_X5.R | TATATGGATCCGCCACCTTAAGCCTCATCAATATGGG | |

48
